# Neuro-Computational Foundations of Moral Preferences

**DOI:** 10.1101/801936

**Authors:** Giuseppe Ugazio, Marcus Grueschow, Rafael Polania, Claus Lamm, Philippe N. Tobler, Christian C. Ruff

**Author notes:** Correspondence to: Giuseppe Ugazio, Geneva Finance Research Institute, University of Geneva, Bd. Du Pont d’Arve 40, 1211 Geneva, CH. Christian Ruff, Department of Economics, Blümlisalpstrasse 10, 8006 Zürich, Switzerland. These authors contributed equally to this paper (Giuseppe Ugazio, Marcus Grueschow).

## Abstract

Moral preferences pervade many aspects of our lives, dictating how we ought to behave, whom we can marry, and even what we eat. Despite their relevance, one fundamental question remains unanswered: Where do individual moral preferences come from? It is often thought that all types of preferences reflect properties of domain-general neural decision mechanisms that employ a common “neural currency” to value choice options in many different contexts. This assumption, however, appears at odds with the observation that many humans consider it intuitively wrong to employ the same scale to compare moral value (e.g., of a human life) with material value (e.g., of money). In this paper, we directly challenge the common-currency hypothesis by comparing the neural mechanisms that represent moral and financial subjective values. In a study combining fMRI with a novel behavioral paradigm, we identify neural representations of the subjective values of human lives or financial payoffs by means of structurally identical computational models. Correlating isomorphic model variables from both domains with brain activity reveals specific patterns of neural activity that selectively represent values in the moral (in the rTPJ) or financial (in the vmPFC) domain. Thus, our findings show that human lives and money are valued in distinct neural currencies, supporting theoretical proposals that human moral behavior is guided by processes that are distinct from those underlying behavior driven by personal material benefit.

## Main Text

Moral preferences play a crucial role in determining how we perceive the world, how we act, and what we like. Differences in moral preferences lie at the heart of many types of conflicts between individuals and groups (1–3) and have even led to wars between nations (4–6). More generally, such differences in moral preferences account for the substantial variation in how we judge the actions of other humans and artificial agents (7). Given the relevance and timeliness of moral preferences, it is remarkable how little we understand about the neural and cognitive mechanisms that determine our moral preferences. Knowledge about these processes is essential for understanding cultural and individual differences in moral perception and behavior (8–10), and for the development of AI choice algorithms that concur with the human understanding of morality (11).

In choice domains other than morality, such as economic decisions, individual preferences have been studied intensely in terms of neural processes that assign values to choice options (12). These values are usually inferred by observing choices and fitting models of the presumed utility derived from characteristics of the choice options (such as their magnitude, price, etc.). Note that this assigned utility differs between individuals and therefore is not identical with representations of the option characteristics themselves (since people differ in how they value these characteristics). While most economic models do not actually assume that utilities are represented cardinally, neuroscientists have nevertheless shown that presumed value signals derived with such models correlate with brain activity (13). For instance, several studies have demonstrated that a person’s economic preferences are reflected in subjective values encoded by activity of the ventral-medial prefrontal cortex (vmPFC), ventral striatum (VS), and posterior cingulate cortex (PCC) (13–16). Based on these findings, it has been proposed that the brain values choice-options on a common scale that may allow us to compare and choose efficiently across many different types of goods. This is thought to hold not only for material goods (e.g., art, food or money) but also for non-material values (e.g., beauty, praise, or status, (17–19)).

In the domain of moral choice, several studies (20–23) have likewise proposed that computing the value of human lives or of human pain may draw on the same neural mechanisms that are involved in computing values of non-moral goods. For instance, the vmPFC has been reported to represent the *expected values* of moral options (i.e., the number of possible deaths multiplied by their respective probability of occurrence) (21). However, as this expected-value computation is objective and therefore identical across different agents, it does not reflect a given individual’s *subjective valuation* of the different options that underlies choices between them. Another study showed differences in the neural correlates of emotional and utilitarian appraisals during moral decisions (20); during utilitarian appraisals, the vmPFC represented the number of people to be saved. Similarly, an earlier study (24) reported that the caudate is involved in representing the utility of the outcomes of a moral choice. These findings, however, do neither reveal if differences in moral preferences result from different sensitivity to these attributes nor if individuals assigning different weights to these attributes would take different moral decisions. Finally, recent studies (22, 25–27) showed that neural value responses in the vmPFC were differentially modulated during choices about financial rewards that were coupled with morally-relevant consequences (e.g. painful shocks to either others or oneself). However, in this context it is impossible to know whether these neural activations indeed reflect moral concerns rather than other aspects of these consequences. For instance, in the case of the pain studies (22, 25), the vmPFC could be responding to differences in the representation of others’ versus one’s own affective states during pain (28, 29). Moreover, since these decisions always entailed trade-offs between pain and monetary profit, the observed neural responses in the value system reflected the valuation of material goods (and how this was altered by different moral contexts). Thus, despite several interesting findings on moral decision making and value computations, we still do not know the neural origins of an individual’s subjective moral preferences and whether the neural mechanisms performing purely moral value computations differ from those involved in the neural valuation of material goods. Clarifying whether moral and material preferences are represented by distinct neural mechanisms is crucial for having a better understanding of decision-making processes that entail a combination of these two types of preferences, such as philanthropy or sustainable finance.

Differences in the neural processes underlying moral and material preferences are suggested by theoretical accounts emphasizing that moral preferences may originate from specific value-computation mechanisms. These accounts rest on the observation that many people perceive human lives as having an intrinsic (sacred) value (30, 31) that cannot, and should not, be measured on the same scale as the value of material objects (32). For example, widespread outrage is usually observed when people realize that the value of human lives is explicitly quantified in terms of money, for instance during choices between health policies (33), in the context of a company’s decision on whether to re-call a dangerous car model (34), or when people are traded for money (35). Based on these observations, it has been proposed that assigning a financial value to a human life appears intuitively wrong for many people (30). This suggests that moral valuation may be implemented by processes that are distinct from those involved in the valuation of material goods.

In the present work, we test this alternative hypothesis by identifying where and how the brain computes purely subjective moral values, and by explicitly comparing the neural instantiation of moral and financial value-computations. We measured these two types of subjective valuation processes with structurally equivalent choice tasks that differed only in the content of the choice-options: Valuing human lives for moral decisions and valuing monetary rewards for financial decisions. We decided to focus on human lives since subjective moral values are essential for the difficult decisions whether some lives are more valuable than others, and since there are considerable individual differences in this regard (7). One example are decisions about recipients of an organ transplant, for which it is often required to implement a policy ranking among the potential recipients to decide who is most deserving to receive the organ (36). We adapted this decision situation to study the neural representations of subjective moral values, which we derived by fitting standard computational models of value-based decision making to the observed choices (37–39) and by correlating the estimated values with neural activity as measured with functional magnetic resonance imaging. More specifically, we used the standard econometric *revealed-preference* approach to estimate the value that each participant assigns to not sacrificing a given person (who is characterized by morally-relevant previous deeds, see the methods below for more details) in order to save a varying, larger group of other people. We did so by applying a standard value-discounting choice model to the observed choices of this participant, which gave us each participant’s subjective value assigned to each specific trial. Note that this subjective value will vary systematically across trials, but also across participants, in line with their moral preference (as derived from the fitted choice model). That is, an individual with a strong moral preference for protecting individual lives will consider it immoral to harm someone even if this can bring about a greater good and will assign a very high subjective value to the life of the person that he may harm. Conversely, an individual with a strong moral preference for bringing about the greater good will assign a low subjective value to the life of the person that will be harmed in order to achieve such greater good. Thus, a given trial/choice problem will elicit very different SVs in participants with different preferences (between-subject variation), and the same participant will assign very different SVs to different choice problems/trials based on the varying moral deservingness and number of people that can be saved on this trial (within-subject variation). By correlating these varying value signals with BOLD signals within subjects, we can thus identify neural signals that reflect individually-specific value computations that are fully in line with each individual’s moral preference, rather than with the objective magnitudes/probabilities/delays characterizing a choice problem (which would not vary across individuals with different preferences, as in e.g. (21, 40)).

In order to fully capture individual behavioral variability during both decision types, we varied the decision-relevant characteristics of the choice options along two dimensions. For the financial decisions, participants chose between options that differed in terms of both the monetary amount and the temporal delay at which the amounts would be paid out. The subjective value of the choice options therefore depends inherently on individual time preferences (15, 41, 42), which determine how the reward magnitude (i.e., the amount of money one can receive) is discounted by the delay (i.e., the number of days) one has to wait until receiving the reward. The moral decisions were constructed to match exactly this structure: They consisted of a customized moral scenario similar to the trolley moral dilemma (43) that required participants to take medical decisions similar to real-life moral choices taken by doctors: They had to choose between a) interrupting life-support of a patient in a coma state to use the patient’s organs to save the lives of other individuals or b) leave the coma-patient on life-support and let the other individuals die (see methods below). For these moral decisions, we parametrically varied both the choice-relevant magnitude (i.e., the number of lives one could save) and a second factor that discounted the value of the lives at stake. This factor was the moral deservingness of the person that would have to be sacrificed in order to save the others (as indicated by different prior criminal records of this person). Both these factors have been shown to play important roles in moral judgments (21, 44), in combination with other factors such as social status and fitness (7). Interestingly, there seem to be fundamental cultural and individual differences in the importance assigned to moral deservingness when people have to estimate and compare the value of human lives (7). This highlights the importance of understanding how individual and situational factors jointly determine moral perception and preferences. Our paper takes an important step in this direction, since it provides a value-computation model that captures moral preferences both behaviorally and in terms of the underlying neural value computations.

Our setup allowed us to directly compare the neural value representations underlying both types of choices in the same participants using functional magnetic resonance imaging (fMRI). We ensured that the perceptual and sensorimotor demands required by both types of choices were kept similar, as the choice screens in both contexts were arranged analogously (Fig. 1a & b) and as responses were given with the same motor actions. Based on the existing value-based literature, we estimated subjective values underlying the financial choices by means of computational modeling (39, 45) and expected to confirm their neural representations in brain activity in the vmPFC, the VS, and the PCC (15, 42, 46). We estimated moral subjective values with structurally isomorphic computational models; this allowed us to test whether moral subjective values would be represented by similar structures as financial values (e.g., the vmPFC, Shenhav & Greene 2010) or whether they instead engage representations in other brain areas (e.g., in the right TPJ, (44, 47), thereby disproving the common currency hypothesis for moral preferences.

**Figure 1:**
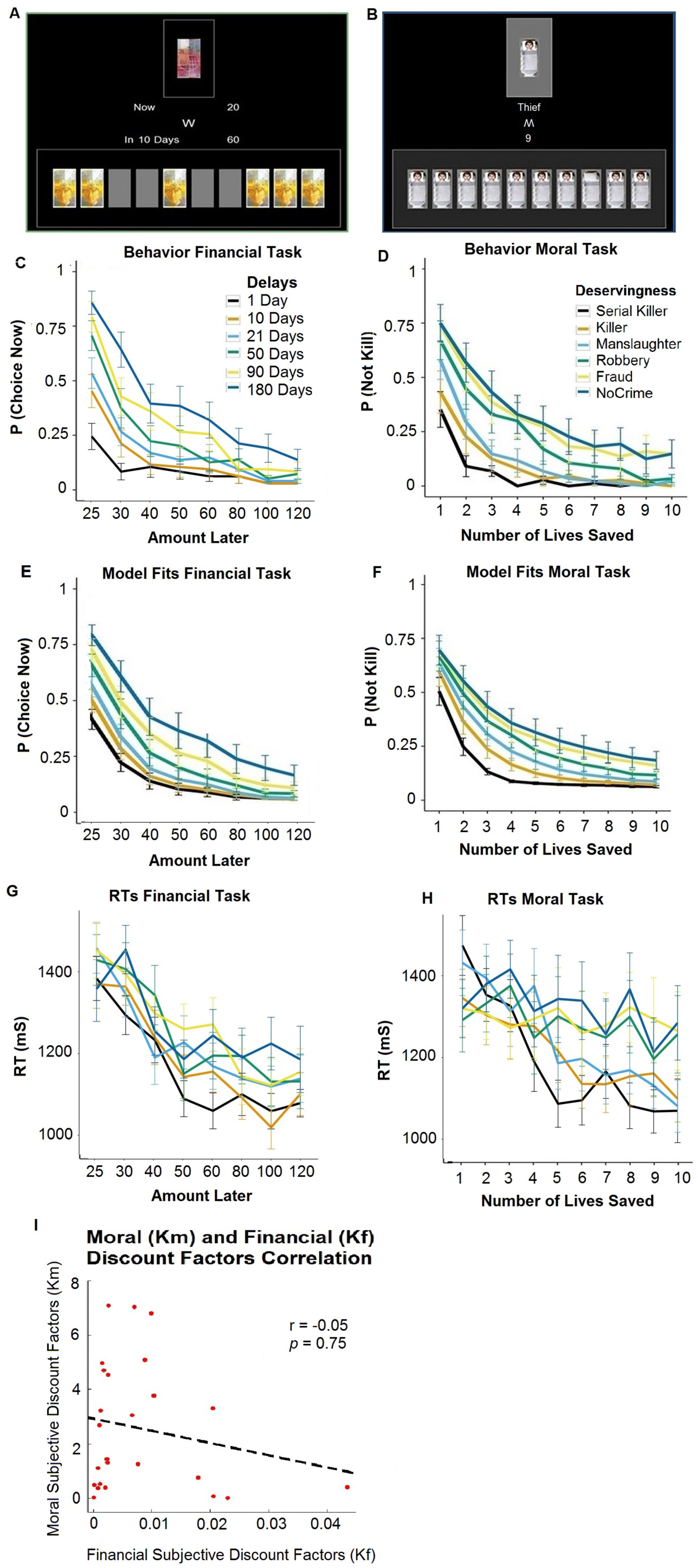
Paradigm and Behavioral Results: Participants made financial (**A**) and moral (**B**) choices. In the financial task, they decided whether or not to give up a sooner smaller financial reward for a later larger financial reward. In the moral task, they decided whether or not to sacrifice one coma-patient to save a larger group of people requiring organ transplants. (**C**) The probability of giving up the sooner-smaller reward increased as the amount of the delayed reward increased. The increase was modulated by the delay participants had to wait to receive the larger option. (**D**) Analogously, the probability of killing the one person in order to save the larger group of people increased with the number of people that could be saved; in this case the probability of choosing to sacrifice the coma-patient was modulated by deservingness. Behavior in both tasks was well captured by the models used, as revealed by the model fits for the financial **(E)** and the moral **(F)** task. Our analyses focused on trials matched for choice (un)certainty, as illustrated by comparable RTs and choice probabilities across the two tasks (**G and H**, analyses of all trials are reported in SI). We found no evidence of correlation between financial and moral discounting (**I**).

## RESULTS

### Behavioral Results

In both types of decisions, participants selected between two choice alternatives on each trial (see the section Materials and Methods in the SI for more information on the tasks): For financial decisions (Fig. 1a), participants chose between 20 Swiss Francs (CHF) to be received today or an equal or larger financial reward (min = 20 CHF, max = 120 CHF) paid out after one of six different time delays (min = 1 day, max = 180 days). For moral decisions (Fig. 1b), participants chose between saving the lives of a larger number of people (min = 1, max = 10) at the expenses of sacrificing the life of one person, or not harming the one person and letting the group die. Moreover, closely mirroring the financial task, participants had to consider an associated feature that may discount the choice option’s value: the moral deservingness of the lives at stake, a property known to play an important role in modulating moral decisions (7, 44). We implemented this by assigning one of six different prior criminal records (ranging from no criminal record to serial killer) to the single person that could be saved or harmed for the benefit of the group. Note that while the two tasks are structurally similar, it is of course possible that different psychological processes may be involved in each of the two tasks. Thus, our setup is not ideal for conducting categorical comparisons across different domains (i.e., if one wanted to test if different brain regions are engaged when people take moral versus financial choices). However, our analyses did not focus on such categorical comparisons but instead investigated correlations of BOLD signals with either moral or financial *subjective values* that varied substantially from one trial to the next within each of the tasks. All these trial-by-trial correlations with subjective values, and comparisons of these correlations between the two tasks, therefore keep constant any factors that differ between moral versus financial decisions per se. Thus, our comparisons of SV representations between the two types of choices cannot be confounded by categorical differences between the psychological processes triggered by the two choice contexts.

Subjective financial and moral values were estimated based on the participants’ financial or moral choices, respectively. In the reward domain, previous studies have repeatedly shown (41, 42) that discount rates are typically well captured by hyperbolic functions, both in humans and other animals (38, 48). In order to estimate participants’ subjective financial values for each trial, we modeled the behavioral data with a standard hyperbolic function:

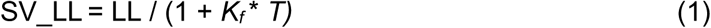

Where *SV_LL* is the subjective financial value of the delayed option estimated as fraction of the immediate reward, LL is the larger later amount offered, *K*_*f*_ corresponds to a subject-specific financial discounting constant, and *T* represents the time (in days) people had to wait to receive the reward. Consistent with previous findings (15), our participants’ discounting curves were well captured by this function (Fig. 1e; R^2^ = 0.98±0.015). Moreover, the financial discount factors (K_f_), and hence the SV_LL, varied substantially across participants (ranging from K_f_ = 3.78*10^−5^ to K_f_ = 0.43, Fig. 1i).

Behavior in the moral task was modeled with a structurally equivalent model to the one used in the financial domain. This allowed us to compare the estimated SV for each task both at the level of behavior (e.g., testing for a correlation between the two SV types) and brain activity. Specifically, behavior in the moral task was modeled with the following hyperbolic function:

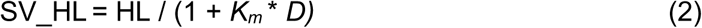

Where *SV_HL* is the trial-wise subjective moral value of saving the lives of the larger group by sacrificing the life of one person; HL reflects the number of human lives one can save in the larger group; *K*_*m*_ corresponds to a subject-specific moral discount factor; and *D* represents the moral deservingness (i.e., criminal record) of the person one could sacrifice. As a first important result, we found that individual discount curves for moral choices (Fig. 1d) were well fit using equation 2 (R^2^ = 0.96±0.03, Fig. 1f). This finding suggests that the moral subjective values estimated here indeed play an important role in moral decision making. Furthermore, like in the financial domain, moral subjective values and the moral discounting factors (K_m_) varied substantially across participants (ranging from K_m_ = 9.3*10^−2^ to K_m_ = 7.08, Fig. 1i).

While the two types of choices were comparable in terms of their computational requirements, they obviously differed qualitatively in terms of choice options and their consequences: On the one hand, participants made decisions about whether or not to harm a human to save other lives, while on the other, they decided between different financial payoffs. It may therefore be expected that the two types of choices may differ in terms of response difficulty. While the average response times (RTs) for the two types of decisions – a standard proxy to measure task difficulty – were similar (average RTs moral 1214ms +/- 28 (s.e.m.), financial 1235ms +/- 24 (s.e.m.), paired t-test, t(24) = 0.39, *p* = 0.7), inspection of the behavioral results (Fig. S1 c and d) revealed a difference in the probability distributions of choosing one of the two options across the two tasks. This difference could indicate that the two tasks may not be fully matched with respect to how Subjective Values relate to choice (un)certainty (49, 50). To control for this potential confound, we focused our SV analyses only on trials that were matched across the two tasks with respect to both response times and the probability of choosing one of the two options. To achieve this, we excluded from the financial task the two trial types that yielded the highest levels of choice certainty (all trials that offered as a larger later reward 20 or 22 CHF).

This exclusion resulted in fully matched choice frequencies and reaction times across both conditions (see Fig. 1 c-d and g-h). We further confirmed this matching in two-sided t-tests comparing the standardized slopes (β1) of a) the logistic regression estimating the probability of choice (see eq.3 and 4 in the behavioral analysis section below) and b) a linear regression (formally, RTs = β0 + β1SV + E) estimating the relation between RTs and moral and financial subjective values for each individual. The t-tests revealed no significant difference across the two tasks, neither with respect to choice probability (t(24) = 0.29 p = 0.77) nor with respect to RTs (t(24) = 0.83 p = 0.41). Thus, following this procedure, the two types of decisions did not differ in terms of choice difficulty, which allowed us to use the model-derived SVs for an unbiased comparison of the underlying neural mechanisms (note that we find similar results when using the complete data-set for this purpose; see supporting information (SI) figures S1-S3 and tables S1, S2).

Interestingly, although financial and moral choices were on average well fitted by identical functions and did not differ with respect to task-difficulty/RTs, we could not find behavioral evidence suggesting that moral and financial valuation processes rely on correlated psychological mechanisms: When testing for a relationship between each individual’s discounting in the financial and moral domain, we found no correlation between both discounting factors Km and K_f_ (r = - 0.05, *p* = 0.75, Spearman regression, Fig. 1i). This absence of a correlation already suggests that moral and financial value estimations may be performed by independent neural and cognitive decision mechanisms.

### Functional Imaging Results

As an initial imaging analysis step, we confirmed the well-known neural correlates of subjective *financial* values. As expected based on the literature (15, 51, 52), we found a significant correlation between subjective financial values of the delayed monetary option (SV_LL) and BOLD activity in brain areas associated with subjective financial value-processing (13, 14). In particular, we found the hypothesized positive financial subjective value representations in the vmPFC and dmPFC (see Fig. 2a and Table 1). We did not find any activation reflecting negative subjective financial values (-SV_LL).

**Table 1:** Average brain activity representing subjective moral values positively (SV_HL, rows 4-7, related to Figure 2B) and negatively (-SV_HL, rows 9-12, related to Figure 2C), and average brain activity representing subjective financial values positively (SV_LL, rows 14-18, related to Figure 2A, no activity was found for -SV_LL). All p-values are FWE-corrected for the whole brain. ACC = anterior cingulate cortex; AntIns = anterior insula; DLPFC = dorsolateral prefrontal cortex; IPL = Inferior parietal lobule;; TPJ = temporo-parietal junction; MPFC = medial prefrontal cortex; SMG = supramarginal gyrus; STS = superior temporal sulcus. Coordinates are listed in MNI space.

| TABLE 1 |  |  |  |  |  |  |  |  |
| --- | --- | --- | --- | --- | --- | --- | --- | --- |
| Region | Peak-Side | Cluster Size | x | y | z | Z score | T score | p-value |
| **Neural Correlates of Subjective Moral Values (SV_HL)** |  |  |  |  |  |  |  |  |
| ACC |  | 839 | 0 | 29 | 40 | 5.14 | 7.06 | <0.001 |
| AntIns | R | 111 | 36 | 2 | 10 | 3.73 | 4.40 | <0.001 |
| IPL | L | 162 | -51 | -37 | 25 | 4.31 | 5.38 | <0.001 |
| IPL | R | 104 | 60 | -16 | 22 | 4.61 | 5.93 | <0.001 |
| **Negative Neural Correlates of Subjective Moral Values (-SV_HL)** |  |  |  |  |  |  |  |  |
| Cuneus | R | 125 | 12 | -85 | 4 | 3.92 | 4.70 | <0.001 |
| DLPFC | R | 829 | 54 | 35 | 22 | 4.80 | 6.30 | <0.001 |
| PCC |  | 506 | 0 | -67 | 37 | 5.21 | 7.22 | <0.001 |
| TPJ | R | 538 | 48 | -58 | 31 | 4.55 | 5.82 | <0.001 |
| **Neural Correlates of Subjective Financial Values (SV_LL)** |  |  |  |  |  |  |  |  |
| MPFC | L | 938 | -12 | 50 | 49 | 5.04 | 6.82 | <0.001 |
| SMG | R | 273 | 66 | -37 | -2 | 4.62 | 5.95 | <0.001 |
| STS | L | 337 | -48 | -61 | 25 | 4.15 | 5.08 | <0.001 |
| STS | R | 135 | 60 | -61 | 28 | 4.28 | 5.32 | <0.001 |
| Visual Cortex | L | 194 | -30 | -52 | -14 | 5.04 | 6.83 | <0.001 |

**Figure 2:**
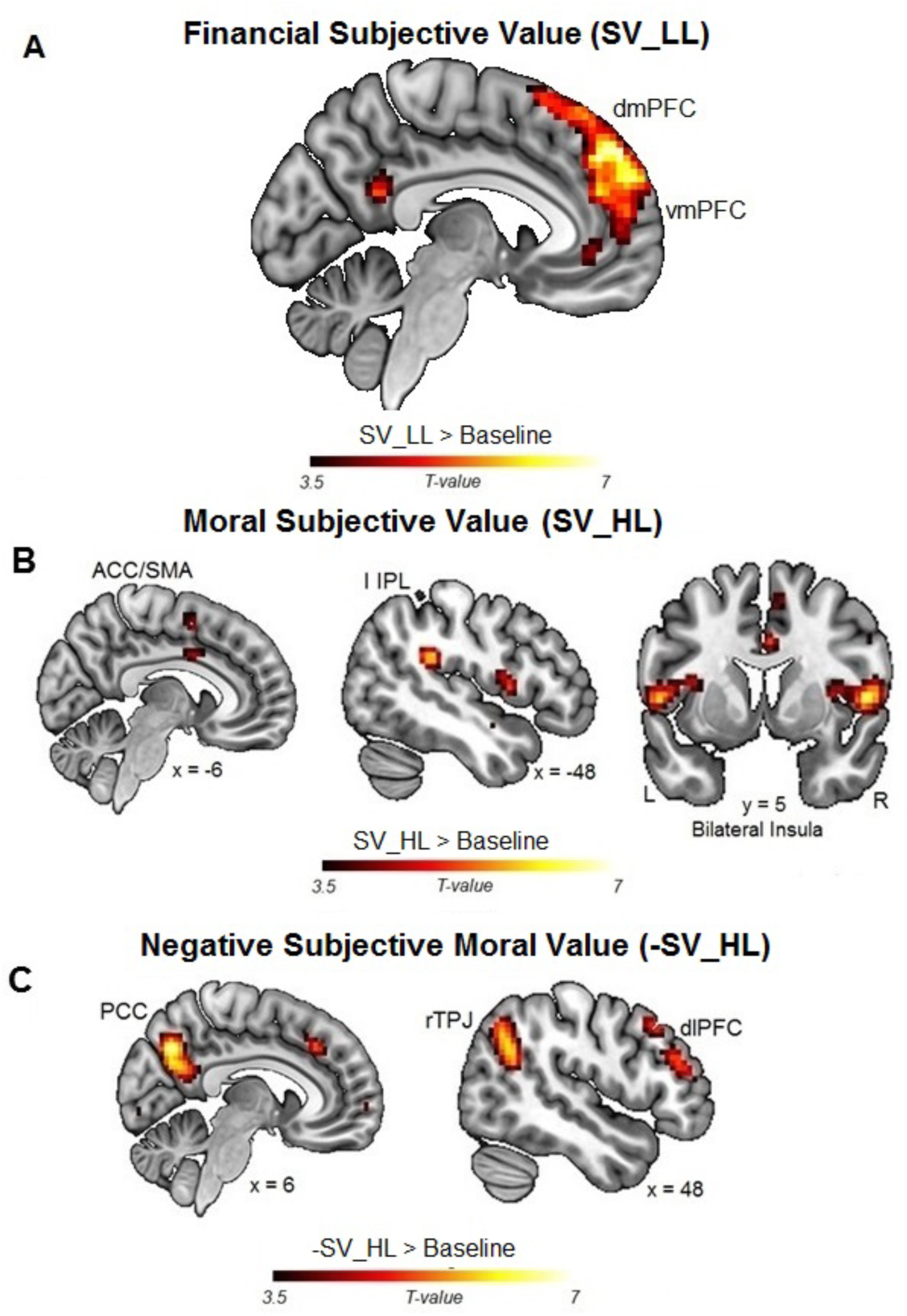
Functional Neuroimaging Results. (**A**) Financial Subjective Values (SV_LL) were represented by neural activity in the mPFC; (**B, C**) Moral Subjective Values (SV_HL) were positively represented by neural activity in the bilateral anterior insula (**B**) and negatively (-SV_HL) in the right TPJ, dlPFC and PCC (**C**)

Importantly, our fMRI analysis revealed that the trial-by-trial subjective *moral* values were represented by BOLD signals in a set of brain regions comprising the bilateral TPJ, the PCC, the right DLPFC, the right anterior insula, the left inferior parietal lobule (IPL), and the anterior cingulate cortex (ACC) (see Fig. 2b and 2c, and Table 1).

These results allowed us to directly relate individual differences in moral preferences to differences in neural activity in these brain regions, effectively providing novel evidence of a neural signature of subjective moral preferences. More specifically, we found that the higher the subjective moral value of the trial-wise varying numbers of human lives at stake (SV_HL), the higher the BOLD activity in the bilateral anterior insula, the left inferior parietal lobule (IPL), and the anterior cingulate cortex (ACC; Fig. 2b), and the lower the BOLD activity in the rTPJ, the PCC and the right DLPFC (Fig. 2b). These results are generally consistent with previous reports of activity in some of these brain areas during moral decisions (20, 44, 53, 54), as well as in the representation of expected values in the moral domain (21), but they now unambiguously reflect subjective moral preferences.

Nevertheless, one may wonder whether our results reflect subjective values representations used to guide choices or may rather be a consequence of these (binary) choices themselves. This is because the tendency to sacrifice the one person to save the group increases with the varying moral subjective value (SV_HL; conversely, the tendency to not sacrifice this person increases with -SV_HL). To investigate this, we ran an additional analysis that, instead of focusing on SVs, identified neural activity correlating with the choice reported by the participants (i.e., moral task: choices not to harm the one person vs. choices to save the larger groups or vice versa; financial task: choices to keep the smaller immediate reward versus the larger later reward or vice versa). These analyses did not reveal significant activations, suggesting that the neural signals identified by the previous analyses indeed reflect subjective value computations and not choice outcomes per se.

A crucial aim of our fMRI analysis was to establish if moral subjective value computations rely on domain-general mechanisms also shared with non-moral value-based decisions (21) or whether they rely on markedly different brain regions. We thus directly compared the neural activity related to SV_HL computations versus the activity related to the matched SV_LL computations. This confirmed that the activity in the rTPJ, the rDLPFC, and the PCC that correlated negatively with SV_HL (i.e., that coded for the moral value of not harming the one person) was indeed domain-specific, as it was significantly stronger than the BOLD correlations with SV_LL (Fig. 3 and Table 2). Intriguingly, the activity in the ACC, insula, and IPS correlating positively with SV_HL (the moral value of saving the larger group) was not domain-specific, i.e., it was not significantly stronger than the BOLD correlations with SV_LL in those regions. In contrast, BOLD activity in the mPFC (Fig. 3 and Table 2) was specifically related to representing financial values, since it was significantly stronger than the BOLD activity correlation with moral subjective value (i.e., SV_LL > -SV_HL).

**Table 2:**
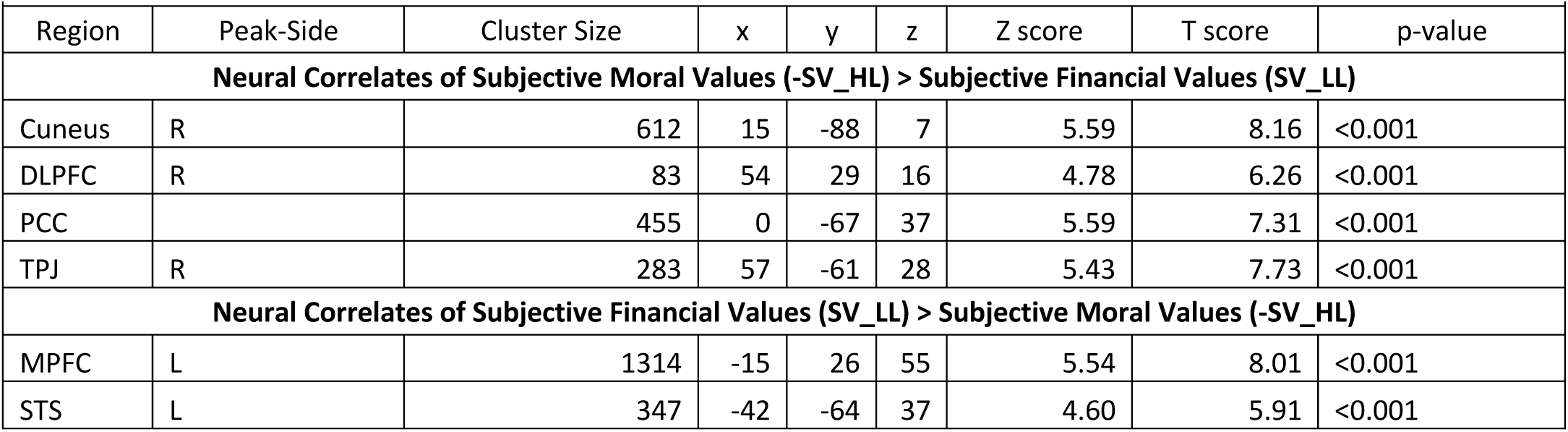
Average brain activity specifically representing subjective moral values > subjective financial values (rows 4-7), and average brain activity specifically representing subjective financial values > subjective moral values (rows 9-10), related to Figure 3. These analyses were performed on trials matched for (un)certainty across the two tasks. All p-values are FWE-corrected for the whole brain. DLPFC = dorsolateral prefrontal cortex; PCC = posterior cingulate cortex; TPJ = temporo-parietal junction; MPFC = medial prefrontal cortex; STS = superior temporal sulcus. Coordinates are listed in MNI space.

| TABLE 2 |  |  |  |  |  |  |  |  |
| --- | --- | --- | --- | --- | --- | --- | --- | --- |
| Region | Peak-Side | Cluster Size | x | y | z | Z score | T score | p-value |
| Neural Correlates of Subjective Moral Values (-SV_HL) > Subjective Financial Values (SV_LL) |  |  |  |  |  |  |  |  |
| Cuneus | R | 612 | 15 | -88 | 7 | 5.59 | 8.16 | <0.001 |
| DLPFC | R | 83 | 54 | 29 | 16 | 4.78 | 6.26 | <0.001 |
| PCC |  | 455 | 0 | -67 | 37 | 5.59 | 7.31 | <0.001 |
| TPJ | R | 283 | 57 | -61 | 28 | 5.43 | 7.73 | <0.001 |
| Neural Correlates of Subjective Financial Values (SV_LL) > Subjective Moral Values (-SV_HL) |  |  |  |  |  |  |  |  |
| MPFC | L | 1314 | -15 | 26 | 55 | 5.54 | 8.01 | <0.001 |
| STS | L | 347 | -42 | -64 | 37 | 4.60 | 5.91 | <0.001 |

**Figure 3:**
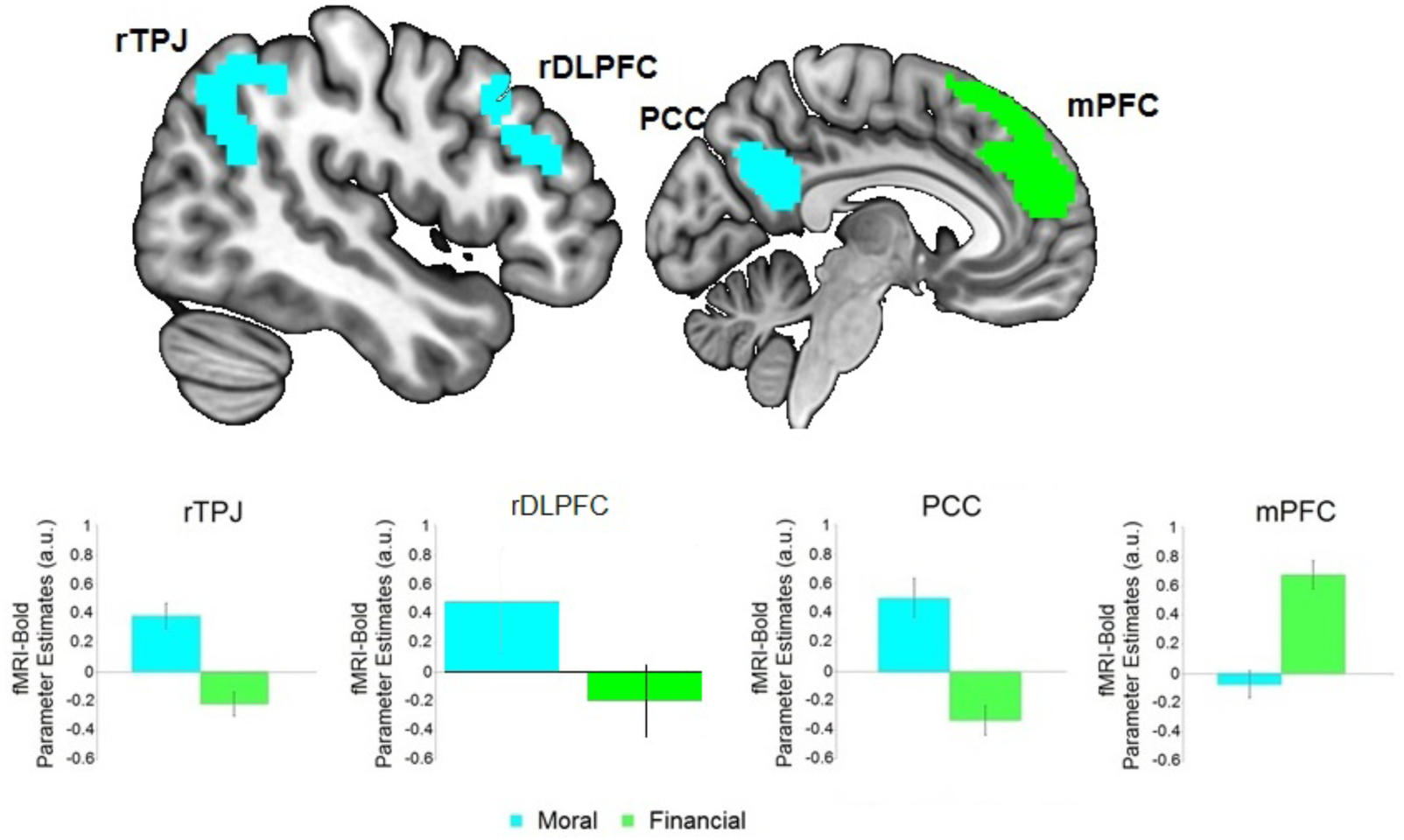
Domain specific subjective value representations in the Financial Task: Specific neural representations of moral subjective value (-SV_HL) were found in the rTPJ, rDPLFC and the PCC (cyan). In contrast, specific correlates of financial subjective value (SV_LL) were identified in the mPFC (green). Colored areas represent clusters of activity specific for each of the two tasks, and not an ROI analysis. These analyses were performed on trials matched on (un)certainty across the two tasks.

To further test if the common neural currency network indeed represented only Financial Subjective Values but not Moral Subjective Values, we performed an ROI analysis (8mm Sphere centered at peak coordinates, see Table S3 in the SI) that tested for neural activity in brain regions indicated by a previous meta-analysis to represent Financial SVs (52). Confirming our whole-brain analysis, this ROI approach also revealed that only Financial Subjective Values were represented within the neural-common-currency network (SV_LL > - SV_HL; t-test, all ps < 0.001, see Table S3 in the SI). These findings highlight that the neural representation of moral subjective values relies largely on domain-specific mechanisms. We also tested for potential functional activity involved in computing both moral and financial subjective values. The conjunction analysis testing for such overlap in coding of both SV_LL and SV_HL, however, did not reveal any significant result.

## Discussion

We identified neural value representations that underlie individual moral preferences, thereby testing if the brain represents moral and material preferences in a common neural currency. This hypothesis would be consistent with widely held views that neural value processes should be domain-general, but contradicts the moral intuition that human lives should not be valued in material terms and in the same currency as objects or cultural artifacts. We tested this hypothesis with a novel moral choice paradigm allowing us to investigate if the human brain explicitly represents the subjective value of saving/harming the life of other persons, and to compare these neural processes with those representing financial subjective value. Our behavioral models show that, similar to financial choices, moral decisions concerning who should be saved/harmed are well fit by computational decision models estimating the subjective value that people assign to the choice alternatives. Neurally, we found that moral subjective values are computed according to similar principles as financial values, but are represented in domain-specific brain areas that differ markedly from those involved in financial valuation.

More specifically, our data show that subjective moral values (+/-SV_HL) are represented in a network of regions comprising the right TPJ, the PCC, the right DLPFC, the left IPL, and in the anterior insula. Importantly, directly comparing the neural activity elicited by moral vs. financial subjective values allowed us to demonstrate that purely moral subjective values are not represented in the same common neural currency representing the value of material types of goods (17–19). Instead, we found that neural activity in the rTPJ, PCC, and other areas (see Fig. 3 and Table 2) was specifically involved in the representation of moral compared to financial subjective values. This suggests that the brain houses a moral-specific valuation system involving neural activity in brain areas previously interpreted as representing socially salient components of decisions, such as empathy (55, 56) harm-aversion (22). Crucially, we found that only neural activity representing the moral value of harming one specific person, captured in our model by -SV_HL, was specifically recruited for moral value-computations; this was not the case for the neural activity correlating positively with the number of lives in the larger group one could save (captured in our model by SV_HL). Thus, our findings support the view that subjective moral valuations recruit both moral-specific valuation mechanisms (in the case of -SV_HL) and but also domain-general decision-processes as previously identified (21). This previous study found that certain choice-relevant information about the options in moral decisions (such as magnitudes and probabilities of outcomes) are represented by domain-general neural mechanisms. In particular, it was found that computations of the expected value of probabilistic outcomes in moral scenarios elicited neural activity in regions commonly associated with computations of the expected value of probabilistic financial rewards, such as the striatum (57) and the vmPFC (16, 58). However, as this study focused on representation and computation of objective information (such as probability, magnitude and expected values), it does not inform us with respect to how the brain represents and derives subjective moral preferences that differ between individuals with different moral stance. Nevertheless, some aspects of our results are consistent with this previous study, since we found that BOLD correlates of the moral value computations taking into account the numbers of lives that one could save in the larger group (i.e., SV_HL) did not differ from those involved in financial choices relying on similar magnitude estimations. Thus, our results show that from the perspective of neural coding, the moral values used for choices comprise both a subjective, domain specific component and a domain-general component shared with financial choices.

Other studies investigating decisions in contexts that require integrating moral and financial values (e.g., deciding between donating to a charity or keeping the money for oneself) found that the subjective value of choice options was mostly represented in the vmPFC but modulated by social information from the TPJ (25, 27, 59, 60). In this decision context, the rTPJ has been thought to estimate socially-salient components, such as the need to overcome one’s perspective or the deservingness of a charity, and to pass this information on to the vmPFC where the value-computation is ultimately implemented (27, 60, 61). However, in all these studies, the choice options resulted in financial payoffs, meaning that they cannot determine whether the vmPFC activity represented financial values (that were modulated by moral concerns) or the moral values themselves. Thus, our results offer a novel and intriguing perspective on the role of the rTPJ in moral value-computations. For decisions based only on moral values (i.e., where there is no trade-off between self-interested financial values and moral values), our data suggest that subjective moral values can be represented directly in the rTPJ without any vmPFC involvement. This suggests that moral preferences originate from the idiosyncratic structural and functional properties of rTPJ (and the other areas we identified) rather than from value coding in the vmPFC.

As noted above, it is of course possible that our two choice tasks differ with respect to the psychological processes involved in the two types of decision contexts. For instance, the two tasks could differ with respect to the amount of imagination/perspective taking involved to make a choice. However, note that any such possible differences are very unlikely to have confounded our results. First, these differences would mainly have affected categorical differences (i.e., comparisons of all moral versus all financial trials), which we did not test for here. Instead, we tested for differences in correlations of BOLD with moral SVs versus with financial SVs. These SVs varied across trials and individuals, so any constant differences between the two types of trials was controlled for in our analyses (it is highly implausible that imagination/perspective-taking should correlate systematically with SVs in just one domain but not the other). Second, it is actually unclear if and to which extent financial and moral decisions differ along these dimensions. Based on previous studies of hypothetical vs. real decisions across different domains (62, 63), one would expect the imagination network to comprise the PCC but also the mPFC and the posterior Hippocampus (pHipp). However, we found that one of these regions (the mPFC) specifically represented financial subjective values, whereas another of these regions (the PCC) specifically represented moral subjective values. Moreover, one previous study (59) demonstrated a causal involvement of perspective-taking processes in the TPJ in classic financial intertemporal choices. Thus, our results are very unlikely to be influenced by general differences in psychological processes triggered by the different choice contexts; they are much more likely to reflect specifically the moral and financial subjective values that varied across trials and individuals.

Relating moral preferences to neural activity, we found that SV_HL was negatively associated with neural activity in the rTPJ, PCC, and DLPFC among other areas (see Table 1 and Fig. 2c), and positively associated with activity in the anterior insula and the left IPL (see Table 1 and Fig. 2B). These findings suggest a mechanistic interpretation of how moral preferences in our choice context are represented in the brain. That is, SV_HL may be computed based on assessments of the harm inflicted on the one person who may be killed as a consequence of one’s choice, consistent with previous studies linking brain activity in the right TPJ, PCC, and DLPFC to processing harm aversion and empathy (22, 64–66). Thus, this neural activity could be interpreted as encoding the increase in value of a human life related to its increasing moral deservingness (which correlates negatively with SV_HL and positively with TPJ/PCC/DLPFC activity). Conversely, the moral preference that may consider it required to save a larger number of people could rely on neural valuation mechanisms responsible for comparing the magnitudes of the moral choice options, reflected in brain activity in the left IPL and in the anterior insula that was not truly domain-specific (i.e., not significantly stronger than the corresponding correlations with the magnitudes of financial values). The left IPL has been associated with magnitude representations and reasoning processes (67), while the anterior insula has been associated with representing social arousal and emotions elicited by socially salient stimuli (68, 69). An increase of anterior insula activation in this case could thus reflect the increased arousal resulting from the increased evidence endorsing a harming action (killing the patient to save the large number of lives; see also (70)). This interpretation accommodates and extends the ideas proposed in the previous study that identified a positive correlation between activity in these brain areas and an increase of expected value in moral decisions with probabilistic outcomes (21). Taken together, our results thus suggest that moral preferences are encoded by (at least) two antagonistic neural systems, rather than by a unitary neural network as is the case for financial preferences.

The analysis of the monetary control task showed that financial subjective values were indeed represented by neural activity in the vmPFC and PCC, consistent with numerous previous findings (15). These value-computation mechanisms may well be domain-general to some degree, and contribute to moral decisions that require representation and integration of objective information about choice outcomes (such as the magnitude and probability of achieving an outcome (21)) or its “utility” to the agent (40). Such domain-general mechanisms for information representation can even be useful to anticipate individual sensitivity to different features of the choice options: For instance, it was shown (21) that participants’ sensitivity to probability of outcomes had higher Ant. Ins. activity, or that participants who were more sensitive to magnitudes displayed higher vmPFC and IPL activity. However, since this study did not identify moral preferences in terms of subjective moral values, it is currently unclear how such these preference computations may relate to, or be influenced by, individual differences in representations of choice options and their characteristics. Nevertheless, these domain-general mechanisms seem to have an important role for moral values when these are traded-off with other types of values, for instance in situations where financial valuation mechanisms corrupt human moral values (71) or where moral values related to the aversion of harming others can discount financial values (25). This raises the interesting question for future studies what context factors may determine how domain-general valuation mechanisms compete and interact with the moral-specific mechanisms identified here, and which higher-level areas may control the interaction of the different valuation systems.

More generally, while trolley-type moral dilemmas have been questioned for their ecological validity (72), recent technological developments in robotics and artificial intelligence have revitalized the importance of studying human decision making in these type of dilemmas (73). Moreover, a previous cross-cultural study (7) demonstrated that while there seem to be general, culture-independent moral preferences (such as saving a larger number of lives), the situational factors influencing moral preferences strongly vary across cultures and countries. In our study, we show that differences among moral preferences can already be detected at the individual level and can be explained using a simple subjective-value computational model. Consistent with the previous behavioral study (7), we found a general preference for saving larger number of lives for all our participants. However, this general preference strongly interacted with each participant’s subjective perception of moral deservingness of the lives involved, and both these factors jointly determined the resulting subjective moral values. Thus, our results provide critical information on the origins of individual and cultural differences in moral preferences, and may be important for future ethical, public and scientific debates regarding decisions taken by artificial intelligence. For instance, it is increasingly debated how a self-driving car should be pre-programmed for selecting whom to harm in potentially critical situations where different lives are at stake - should it always protect the people inside the car or should it use some other criterion? Our current results identify distinct neural mechanisms by which our brains compute tradeoffs between saving and harming human lives, which differ from neural valuation processes involved in selecting between material goods. This suggests that artificial intelligence may benefit from accounting for the properties of these mechanisms in order to be perceived as morally appropriate. Last but not least, our study illustrates how moral preferences may be assessed in a manner that is computationally similar to the assessment of financial preferences, without requiring the participants to read and understand complex moral vignettes. This facilitates identification of the choice-related brain mechanisms and may prove essential for moving towards an integrated perspective of how the brain controls and integrates moral and material concerns in the control of actions, in particular in situations where these two types of concern may compete or interact (e.g. in philanthropy or sustainable finance).

## ACKNOWLEDGEMENTS

We thank Karl Treiber for scanning assistance. C.C.R. received funding from the European Research Council (ERC) under the European Union’s Horizon 2020 research and innovation programme (grant agreement No 725355; ERC Consolidator grant BRAINCODES). R.P. received funding from the ERC (grant agreement No 758604; ERC starting grant (ENTRAINER). This work was also funded by grants of the Swiss National Science Foundation (PBZHP1_147240, PP00P1_128574, PP00P1_150739, 100014_165884, 105314_152891, 320030_143443, and CRSII3_141965) to G.U., P.N.T, and C.C.R. All authors gratefully acknowledge support by the Zurich Center for Neurosciences (ZNZ). Author contributions: G.U. and C.C.R. conceived the study; G.U., M.G., C.L., P.N.T., and C.C.R. designed the study; G. U., performed experiments; G. U., M.G., and R.P. analyzed data with conceptual input from C.L., P.N.T., and C.C.R.; G. U., M.G., R.P., C.L., P.N.T., and C.C.R. wrote the manuscript.

