## Supplemental Information for "Neuro-Computational Foundations of Moral Preferences"

* Correspondence to:

**This file includes:**

Materials and Methods

Behavioral Results

Neuroimaging Results

Figs. S1 to S4

Tables S1 to S3

**Materials and Methods**

**Experimental procedures**

**Participants**

The participants were twenty-five healthy students from the University of Zurich (age: min 19, max 34, mean = 22.08, S.E.M. = 0.74 years old; 13 females) with no reported history of neurological or psychiatric disorder and no current use of medication as measured with standard surveys. All the experimental procedures were approved by the Research Ethics Committee of the Canton of Zurich.

**Experimental Task**

Participants made financial and moral choices in randomly alternating blocks during the event-related fMRI sessions. Visual stimulation was highly similar (cf. Fig. 1a and 1b) while the required motor commands where identical. Both tasks were cued visually (a ‘W’ cued financial trials, while moral trial were cued by the letter ‘M’). We chose to indicate the monetary condition with the letter ‘W’ for two reasons: First, it is the first letter of the German word “wirtschaftlich” (“economic”; our participants were students of the University of Zurich and all spoke German fluently). Second, it is visually very similar (inverted) to the “M” used for the moral condition.

During financial choice trials, participants indicated whether they preferred to give up a reward of 20 CHF that was paid out immediately after the study in order to receive a variable reward (Magnitude, 10 levels: 20, 22, 25, 30, 40, 50, 60, 80, 100 or 120 CHF) after waiting different amounts of days (Delay, 6 levels: 1, 10, 21, 50, 90, or 180 days). Thus, the financial task comprised 60 unique combinations of magnitude and delays, which were presented to participants four times – one per each choice-task-session, yielding a total of 240 trials (see SI for the analyses including all trials). After the trial-exclusion procedure to match (un)certainty across the two tasks (see Results section above and in the SI), 48 trials were excluded (2 levels of magnitude combined with 6 levels of delays in each of the four choice-task-session), leaving a total of 192 trials in the main analysis. These financial decisions were fully incentivized: one financial trial was randomly selected at the end of the study and paid out to the participant as described above. If a delayed option was selected, the money was sent via mail to the address specified by the participant. The moral task required participants to read a moral scenario before starting the fMRI session. This moral scenario instructed them to think like a doctor who is taking care of a different patient in a coma-state every day. The moral deservingness of these patients differed as indicated by different criminal records (Deservingness, 6 levels: no records of criminal activities, fraud, robbery, manslaughter, killing a person, killing multiple persons). While on duty, this doctor is informed about the sudden need of organ transplants in variable amounts of victims due to an accident (Magnitude, 10 levels: from a minimum of 1 to a maximum of 10 victims). These people would die if they did not receive the organs soon.

The participant is then asked what she/he would be morally required to do if she/he were the doctor. Choice alternative 1: interrupt life-support to the coma patient, resulting in his death, and use the organs to save the lives of the accident victims; choice alternative 2: leave the coma-patient on life-support and let the victims of the accident die. During the moral choice trials, participants reported which course of action was morally required. Each trial consisted of a unique combination of deservingness and number of people the doctor could save by harming the coma-patient. As for the financial Task, the moral task also consisted of 60 unique combinations of magnitude and deservingness, which were presented to participants four times – one per each choice-task-session, yielding a total of 240 trials (no trials had to be excluded). Critically, our moral task did not require participants to read lengthy and complex moral vignettes before reporting their decisions, but instead presented simple binary decisions where the only varying elements influencing the decisions were the two experimental variables magnitude and deservingness. Note that moral decisions were - for obvious reasons - hypothetical decisions with no practical consequences.

**Behavioral analysis**

Reaction times were analyzed with a two-sided paired t-test comparing the individual average RTs in the moral compared to the financial task.

Financial and moral choices were estimated using structurally identical models of subjective value. Financial choices were estimated with the logistic choice model:

$P_{\mathrm{choice}}= \frac{1}{1+exp(-(B_{0}+B_{1}\times[20-SV\_LL\left( kf \right)]}$	(3)

where the function SV_LL*(k_f_)* is the monetary subjective value in the financial choice (defined in Eq. 1, see Results section) and the free parameter *k_f_* corresponds to the discount factor of the hyperbolic function. The two other free parameters B_0_ and B_1_ were the choice bias and the temperature parameter the logistic regression, respectively.

Moral choices were estimated with a structurally isomorphic model:

$P_{\mathrm{choice}}= \frac{1}{1+\exp(-(B_{0}+B_{1}\times[1-SV\_HL\left( km \right)]}$	(4)

where the function SV_HL *(k_m_)* is now the subjective moral value in the moral choice (defined in Eq. 2, see Results section) and the free parameter *k_m_* corresponds to the discount factor of the hyperbolic function. The other two free parameters are the same as for eq.3.

In both cases, the fitting strategy was based on a Bayesian Hierarchical Modeling (BHM) approach. This approach constitutes an attractive compromise between the extremes of complete pooling and complete independence (1). As in the complete independence approach, BHM estimates parameters for each individual participant. However, these estimates avoid the averaging artifacts that come with the complete pooling approach as well as the unreliability that comes with the estimation of parameters for individual participants. A Bayesian model was fit for each choice type (moral and financial) and contained random effects for the three subject-specific parameters (B_0_, B_1_, and the subjective discounting factors *k_f/m_*) of interest, assuming flat priors for all parameters at the highest hierarchy level. Inference of the parameters in the BHM was performed via the Gibbs sampler using the Markov Chain Montecarlo (MCMC) technique implemented in JAGS (2, 3). A total of 10,000 samples were drawn from an initial burn-in step and subsequently a total of 10,000 new samples were drawn with three chains (each chain was derived based on a different random number generator engine, and each with a different seed). We applied a thinning of 10 to this final sample, thus resulting in a final set of 1,000 samples for each parameter. This thinning assured that the final samples were auto-decorrelated for all of the latent variables of interest. We conducted Gelman–Rubin tests for each parameter to confirm convergence of the chains. All latent variables in our Bayesian models had $\hat{R}<1.05$, which suggests that all three chains converged to a target posterior distribution.

**fMRI data-acquisition and pre-processing.**

Subjects performed four choice-task-sessions (each containing 60 financial and moral perceptual choices) and one resting-state-session that lasted 6.5 minutes each. During each session, we acquired 270 T2*-weighted whole-brain echo planar images using a Philips Achieva 3 T whole-body scanner (Philips Medical Systems, Best, The Netherlands) equipped with an 8-channel Philips sensitivity-encoded (SENSE) head coil. Imaging parameters were: 2600 ms repetition time (TR); 37 slices (transversal, ascending acquisition); 2.6 mm slice thickness; 2.5 mm x 2.5 mm in-plane resolution; 0.65 mm gap; 90° flip angle. To measure at fully equilibrated magnetic field, five dummy image excitations were performed and discarded before functional image acquisition started. To enhance BOLD-contrast sensitivity throughout the brain, we used a dual-echo-sequence (TE: 17 ms and 44 ms) in combination with a weighted voxel-wise summation technique (4, 5) that generates a single functional whole-brain image with optimal sensitivity for each TR. For this procedure, the signal-to-noise ratio is first computed for each echo image voxel in the resting-state scan. These SNR measures are then used to weight each voxel in the two echo images acquired per TR of the choice-task sessions according to the formula

$$X=\frac{X_{E1} \cdot{SNR}_{E1} + X_{E2} \cdot{SNR}_{E2}}{{SNR}_{E1} + {SNR}_{E2}}$$

where X is the resulting image for a given TR, X_E1_ and X_E2_ are the images acquired at that TR for the first echo and second echo, respectively, and SNR_E1_ and SNR_E2_ are the signal-to-noise images (generated as voxel-wise mean divided by the voxel-wise standard deviation) for the resting-state time-series acquired for the first echo and second echo, respectively. A high-resolution T1-weighted whole brain structural image used for image registration during post-processing (181 sagittal slices; matrix size: 256 x 256; voxel size: 1 x 1 x 1 mm ; TR/TE/TI: 8.3/2.26/181 ms) was also acquired for each subject.

Image preprocessing and analysis were conducted using SPM8 (Wellcome Trust Centre for Neuroimaging). Functional images were slice-time corrected (to the middle slice acquisition time) and realigned (accounting for individual head motion). Each participant’s T1-weighted structural image was co-registered with the mean functional image and normalized to the standard T1 MNI template using the new-segment-procedure provided by SPM8 (6). The functional images were then normalized to the standard MNI template using the same transformation, spatially resampled to 3 mm isotropic voxels, and smoothed using a Gaussian kernel (FWHM, 8mm).

**fMRI data-analysis**

The general linear model (GLM) we implemented was suited to identify and contrast correlations of BOLD signals with financial and moral subjective values during the financial and the moral trials respectively. The included regressors were therefore (column 1) an indicator function for financial choices with (columns 2+3) the parametric modulators for trial-wise changing delays and financial subjective values (z-scored at the level of each participant), 4) an indicator function for moral choices with (columns 5+6) the parametric modulators for trial-wise changing deservingness and moral subjective values (z-scored at the level of each participant). The two parametric modulators for financial and moral subjective values were orthogonalized with respect to the parametric modulators for delays and deservingness, respectively, to identify the brain regions in which neural activity is related only to the unique variance of SVs independent of delay/deservingness. That is, our analysis explicitly controlled for potential effects of the objective delay and deservingness on the neural activity elicited by financial and moral subjective values, respectively, by accounting for the corresponding variance. In a control analysis, we tested for neural correlates of the particular choices taken by participants. The analysis included the following regressors: For the financial task 1) trials in which participants chose the smaller immediate option, 2) trials in which participants chose the larger later option. For the moral task 3) trials in which participants chose to not harm the one person, and 4) trials in which participants chose to save the larger group by sacrificing the one person. In addition to these main regressors, all GLMs also included several regressors of no interest: two indicator functions for financial and moral block cues, and six motion parameters (obtained during the realignment procedure). Apart from the motion parameters, all regressors were modelled as stick-functions at the time of stimulus onset, convolved with a canonical hemodynamic response function. The blood oxygen level-dependent (BOLD) signal in each voxel was then regressed onto the composite model using the procedures implemented in SPM8.

First-level summary statistics were obtained by calculating single-subject voxel-wise contrasts for each of the two subjective value parametric modulators. Second-level random-effects group contrast maps were tested for significance by one-sample t tests across single-subject contrast maps. Statistical inference was performed at the cluster level, using a whole-brain FWE-corrected statistical threshold of P < 0.05 (based on a cluster-forming voxel cut-off set to P < 0.001). While this inference procedure provides adequate control of false-positive rates (7), we nevertheless ensured the robustness of our results by replicating the analyses using non-parametric tests (implemented in SnPM). All significant results were confirmed by this second analysis.

**Behavioral Results**

Inspection of behavioral results (Fig. S1 c and d) revealed a difference in probability distributions of choosing one of the two options across the two tasks: in the Financial Task, the probability distribution of choosing the option spans from 1 to 0, whereas in the Moral Task, the probability of harming a human life only spans from 0.77 to 0. This difference could indicate that the two tasks were not matched for choice (un)certainty, in particular on those trials with higher probability of choosing the immediate option in the Financial Task, and that thus the differences in neural representations of the two types of SVs may be reflecting this difference (8, 9). We therefore matched the two types of choices in terms of choice (un)certainty by removing the trials with later payoffs of 20 and 22 CHF form the financial task; in the main text, we report only the SVs analyses focused on these (un)certainty-matched trials (see trial selection procedure on p.9 and Fig. 1). For transparency and completeness, we illustrate the analyses including all trials here (Figure S1).

**fMRI Results**

We repeated the same GLM analyses described in the main paper (see Methods, fMRI Analyses), but this time including all trials to determine the robustness of our results. These analyses yielded comparable results to those reported in the main text. However, in addition to the vmPFC and the dmPFC, we found that also the PCC (FWE-corrected for the whole brain, p < 0.001) and the VS (small-volume-corrected P < 0.05, see Fig. S2a and Table S1) correlated with financial subjective values. These results are largely consistent with previous studies (10, 11). The additional activations found here may result from an increase in precision of parameter estimation, since this analysis included a larger number of trials for the Financial Task. The neural representations of Moral SVs were identical between the two types of analyses (see Table S1 below and Figure S2 B and C below).

Crucially, the analysis investigating specific neural representations of Moral and Financial SVs also yielded comparable results (see Table S2 and Figure S3 below). In this case the only difference is that analysis of all trials no longer yields a significantly stronger representation of Moral SV in the rDLPFC (the p-value rises above the threshold).

Finally, we performed one additional analysis on the entire data-set to investigate if we could detect differences in Financial and Moral SVs representations in an ROI analysis (8mm Sphere centered at peak coordinates, see Table S3 below) that tested for neural activity in brain regions indicated by a previous meta-analysis to represent several Financial SVs (11). Confirming our whole-brain analysis, this ROI approach also revealed that only Financial Subjective Values were represented within the neural-common-currency network (t-test, all *p*s < 0.001).

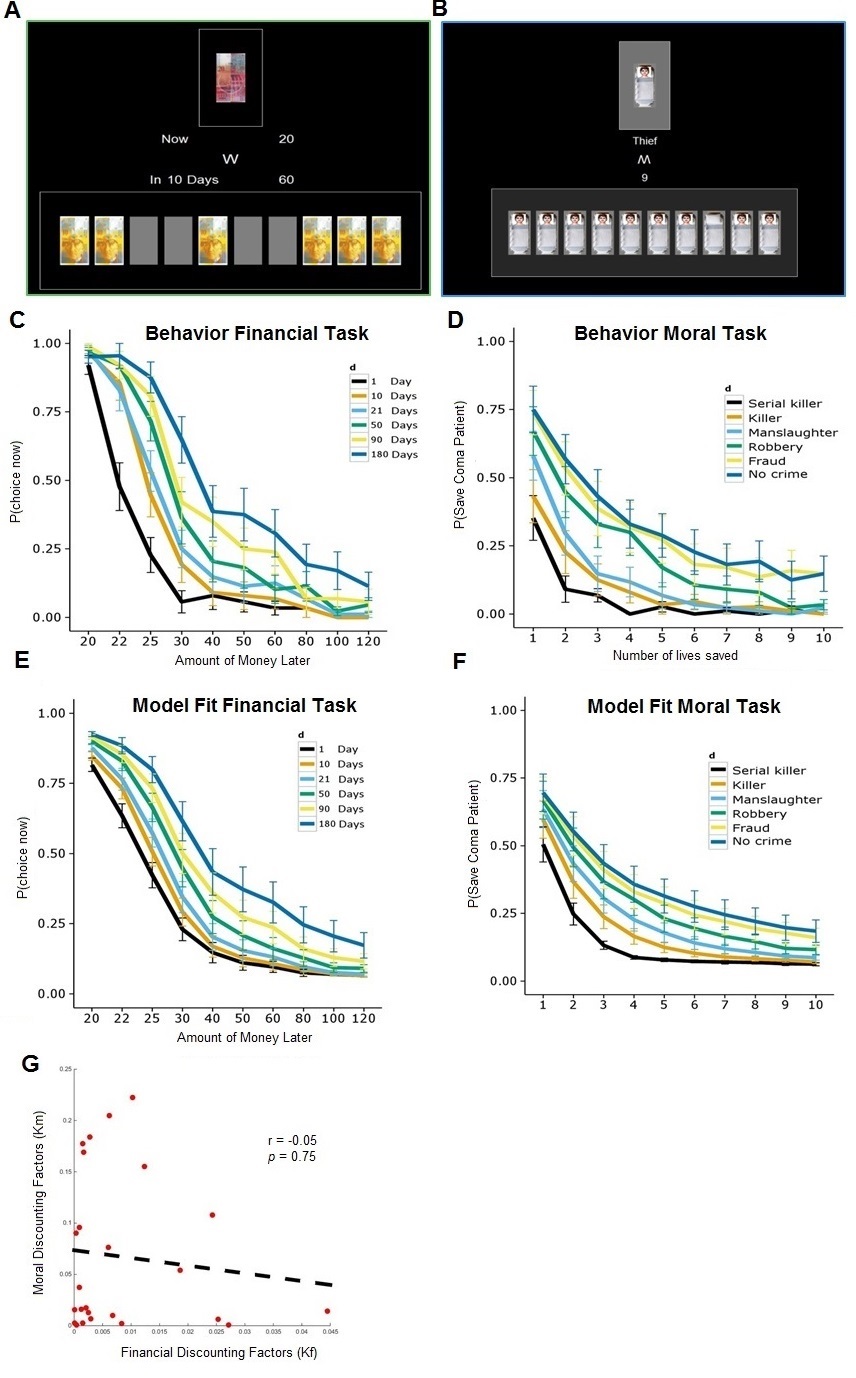

**Figure S1:** Paradigm and Behavioral Results: Participants made financial (**A**) and moral (**B**) choices. In the financial task, they decided whether or not to give up a sooner smaller financial reward for a later larger financial reward. In the moral task, they decided whether or not to sacrifice one coma-patient to save a larger group of people requiring organ transplants. (**C**) The probability of giving up the sooner-smaller reward increased as the amount of the delayed reward increased. The increase was modulated by the delay participants had to wait to receive the larger option. (**D**) Analogously, the probability of killing the one person in order to save the larger group of people increased with the number of people that could be saved. This increase in choosing to sacrifice the coma-patient was modulated by deservingness. Behavior in both tasks was well captured by the models used, as revealed by the model fits for the financial **(E)** and the moral **(F)** task. We found no evidence of correlation between financial and moral discounting (**G**).

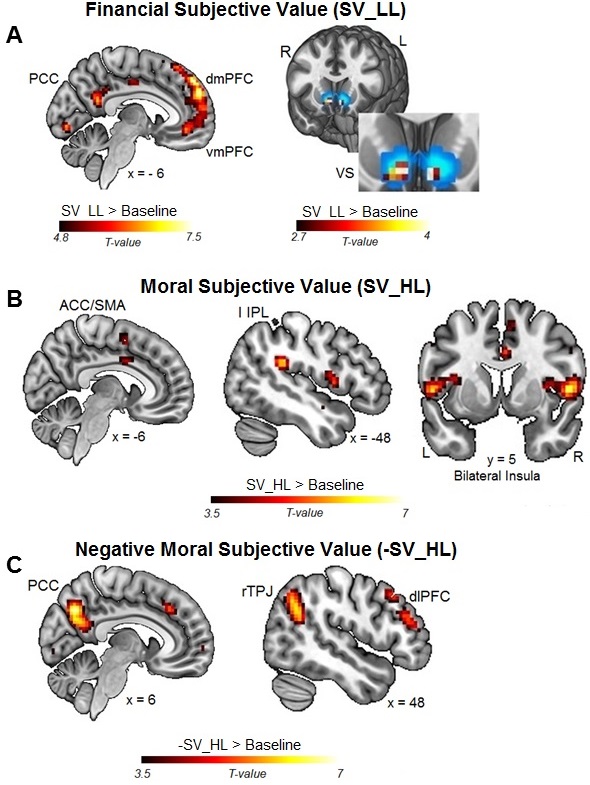

**Figure S2:** Functional Imaging Results. (**A**) Financial subjective values (SV_LL) were represented by neural activity in the mPFC, the PCC, and the VS, consistent with previous findings; Cyan area (right) corresponds to the nucleus accumbens volume mask provided by the FSL-Harvard-Oxford-atlas. (**B, C**) Moral subjective values (SV_HL) were positively represented by neural activity in the bilateral anterior insula (**B**) and negatively in the right TPJ, DLPFC, and PCC (**C**).

| **TABLE S1** | | | | | | | | |
| --- | --- | --- | --- | --- | --- | --- | --- | --- |
| Region | Peak-Side | Cluster Size | x | y | z | Z score | T score | p-value |
| **Neural Correlates of Subjective Moral Values (SV_HL)** | | | | | | | | |
| ACC |  | 839 | 0 | 29 | 40 | 5.14 | 7.06 | <0.001 |
| AntIns | R | 164 | 36 | 2 | 10 | 3.78 | 4.48 | <0.001 |
| IPL | L | 105 | -51 | -37 | 25 | 4.62 | 5.94 | <0.001 |
| IPL | R | 110 | 60 | -16 | 22 | 4.28 | 5.32 | <0.001 |
| **Negative Neural Correlates of Subjective Moral Values (-SV_HL)** | | | | | | | | |
| Cuneus | R | 150 | 15 | -88 | 4 | 4 | 4.83 | <0.001 |
| DLPFC | R | 839 | 48 | 23 | 43 | 4.86 | 6.44 | <0.001 |
| PCC |  | 489 | 0 | -67 | 37 | 5.22 | 7.23 | <0.001 |
| TPJ | R | 518 | 48 | -58 | 31 | 4.59 | 5.89 | <0.001 |
| **Neural Correlates of Subjective Financial Values (SV_LL)** | | | | | | | | |
| MPFC | L | 1831 | -9 | 50 | 43 | 6.39 | 10.57 | <0.001 |
| MTG | L | 238 | -57 | -7 | -14 | 4.44 | 5.6 | <0.001 |
| PCC |  | 194 | 0 | -49 | 31 | 4.69 | 6.08 | <0.001 |
| SMG | R | 273 | 63 | -25 | 1 | 4.34 | 5.42 | <0.001 |
| STS | L | 597 | -57 | -37 | 25 | 4.53 | 5.74 | <0.001 |
| STS | R | 139 | 45 | -28 | 22 | 3.64 | 4.28 | <0.001 |
| Visual Cortex | R | 1777 | 12 | -85 | 4 | 5.87 | 8.92 | <0.001 |

**Table S1: Average brain activity representing subjective moral values positively (SV_HL, rows 4-7, related to Figure S2B) and negatively (-SV_HL, rows 9-12, related to Figure S2C), and average brain activity representing subjective financial values (SV_LL, rows 14-18, related to Figure S2A) in (un)certainty matched trials.**

All p-values are FWE-corrected for the whole brain. ACC = anterior cingulate cortex; AntIns = Anterior Insula; DLPFC = dorsolateral prefrontal cortex; IPL = Inferior parietal lobule; PCC = posterior cingulate cortex; TPJ = temporo-parietal junction; MPFC = medial prefrontal cortex; MTG = medial temporal gyrus ; PCC = Posterior Cingulate Cortex; SMG = supramarginal gyrus ;STS = superior temporal sulcus. Coordinates are listed in MNI space.

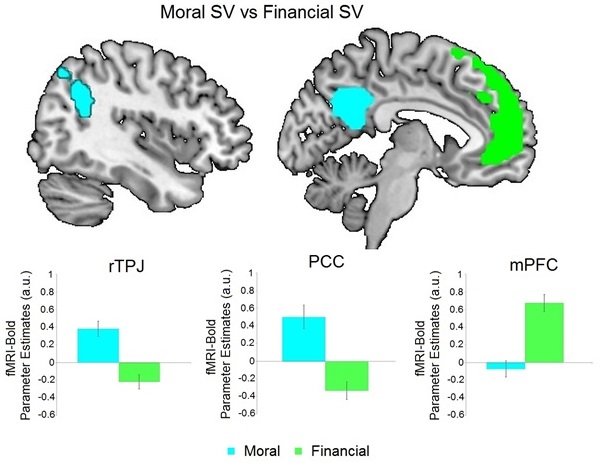

**Figure S3:** Domain specific subjective value representations in the Financial Task: Specific neural representations of moral subjective value (-SV_HL) were found in the rTPJ, rDPLFC and the PCC (cyan). In contrast, specific correlates of financial subjective value (SV_LL) were identified in the mPFC (green). Colored areas represent clusters of activity specific for each of the two tasks, and not an ROI analysis.

| **TABLE S2** | | | | | | | | |
| --- | --- | --- | --- | --- | --- | --- | --- | --- |
| Region | Peak-Side | Cluster Size | x | y | z | Z score | T score | p-value |
| **Neural Correlates of Subjective Moral Values (-SV_HL) > Subjective Financial Values (SV_LL)** | | | | | | | | |
| Cuneus | R | 1908 | 15 | -88 | 4 | 6.35 | 10.44 | <0.001 |
| PCC |  | 471 | 0 | -55 | 25 | 5.25 | 7.31 | <0.001 |
| TPJ | R | 283 | 57 | -64 | 28 | 5.02 | 6.78 | <0.001 |
| **Neural Correlates of Subjective Financial Values (SV_LL) > Subjective Moral Values (-SV_HL)** | | | | | | | | |
| MPFC | L | 1303 | -15 | 47 | 40 | 5.56 | 8.07 | <0.001 |
| STS | L | 347 | -36 | -61 | 25 | 5.16 | 7.09 | <0.001 |

**Table S2: Average brain activity specifically representing subjective moral values > subjective financial values (rows 4-6), and average brain activity specifically representing subjective financial values > subjective moral values (rows 8-9), related to Figure S3.**

All p-values are FWE-corrected for the whole brain. PCC = posterior cingulate cortex; TPJ = temporo-parietal junction; MPFC = medial prefrontal cortex; STS = superior temporal sulcus. Coordinates are listed in MNI space.

| **Table S3** | | | | | | | | |
| --- | --- | --- | --- | --- | --- | --- | --- | --- |
|  | ROI Coordinates (MNI) | | | Peak Activity Coordinates  (MNI) | | | | |
|  | X | Y | Z | **SV_LL** | X | Y | Z | **SV_HL** |
| **mPFC** | 2 | 48 | -8 | P < .001 | -3 | 44 | -11 | - |
| **PCC** | -4 | -30 | 36 | P < .001 | -3 | 37 | 36 | - |
| **R Striatum** | -12 | 12 | -6 | P < .001 | -6 | 14 | -8 | - |
| **L Striatum** | 12 | 10 | -6 | P < .001 | 9 | 14 | -8 | - |

**Table S3: Average brain activity specifically representing subjective financial values or moral values respectively, within the SV ROIs from Bartra et al., (2013)**

All p-values are FWE-corrected for each ROI determined as an 8mm sphere centered at the peak coordinates reported in Bartra etl a., (2013) for each brain reigion: PCC = posterior cingulate cortex; MPFC = medial prefrontal cortex; Coordinates are listed in MNI space.
